# Interplay of microRNA-20a and HuR in the regulation of beclin1 during Withaferin-A mediated impaired autophagy in Breast Cancer cell-line, MCF-7

**DOI:** 10.64898/2026.01.13.699223

**Authors:** Soumasree De, Somdatta Ghosh, Sayantani Das, Alina Chakraborty, Sumita Sengupta (Bandyopadhyay)

## Abstract

Dysfunctional autophagy is connected to multiple diseases. Withaferin-A, a biologically active withanolide, has been shown to impair autophagy in the breast cancer cell line, MCF-7; however, the underlying mechanism remains unclear. Here while uncovering the mechanism, it was found that treatment of MCF-7 cells with WA declines Beclin1 protein synthesis by restricting association of its mRNA to polysomes with no significant reduction in its mRNA level. Reporter assay established that WA-treatment enhanced the expression level of mature miR-20a which directly interacts with the 3’UTR of beclin1 mRNA. RNA affinity chromatography revealed association of mRNA stabilizing protein, HuR with the 3’UTR of beclin1 mRNA. Additionally, co-immunoprecipitation assay determined the interactions of GW182, an essential component of GW-body and Ago2, a crucial member of miRNA related silencing complex (RISC) with the 3’UTR of beclin1 mRNA. In parallel, it was found that knocking down of HuR eliminates the interaction of GW182 with beclin1 3’UTR, while the association of miRNA (Ago2) remains unaffected. These results suggested simultaneous association of HuR, GW182 and microRNA with beclin1 mRNA which could be pivotal for sequestration of beclin1 mRNA in GW-bodies. Thus, our findings culminated in identifying an unconventional mechanism of regulation of Beclin1 expression by the dual interplay between hsa-miR-20a and HuR during WA-induced impaired autophagy in MCF-7 cells.

## Introduction

Autophagy is a complex cellular process that contributes to cellular homeostasis by recycling the essential cytoplasmic components and disposing excessive or defective organelles of the cells [1, 2]. The entire process of autophagy is orchestrated by a well-balanced network of more than 30 major genes (named as *atg*s) [3, 4]. Defects in autophagy can cause different diseases like cancer, aging, infections, neurodegenerative diseases and metabolic disorders [5–8]. Accumulation of autophagosomes and lysosomes by deregulated expression of *atg* genes lead to non-functional cellular recycling machinery causing incomplete autophagy and cellular death [9–12].

Ghosh et al., 2017 had established that Withaferin-A (WA), an active withanolide derived from *withania somnifera,* caused impaired autophagy in breast cancer cell-lines MCF-7 and MDA-MB-231 [13]. WA hampered expression of some *atg* genes, including *beclin1* and also caused defect in autophagosome-lysosome fusion process. Beclin1 is one of most studied effector protein and reported to be essential in autophagosome nucleation and maturation [14]. Mono allelic deletion of Beclin1 is associated with breast, ovarian and prostate cancer. Under normal condition, Beclin1 protein stays associated with anti-apoptotic Bcl-2 family of proteins, through BH3 domain preventing induction of autophagy and apoptosis [15]. Phosphorylation of either BECN1 by DAPK or Bcl2 by JNK disrupts their interactions, thus releases adequate amount of BECN1 necessary for induction of autophagy. BECN1 can offer implications in cancer therapeutics by acting both as tumor suppressor (initiating autophagy [16] or tumor promoter [17]. At this point, understanding the complex molecular mechanism that regulates the expression of this important protein in a context dependent manner became very intriguing.

Complex interplay of the RNA-Binding-Proteins (RBPs) and microRNAs (miRNAs) with the target mRNA can decide its fate at post-transcriptional level [18–19]. RBPs and miRNAs are the major *trans*-factors that can interact with specific *cis*-elements on the 3’-UTR of the target mRNAs [20–21] and thus can modulate mRNA stability, turnover and translation [22–24]. These kinds of RBPs-miRNAs interactions are diverse with different types of diseases or variables with different physiological conditions like hypoxia, stress *etc*. [25–26].

*Trans*-acting proteins include, HuR, nucleolin, PCBP *etc*. act as stabilizing factor, whereas TTP, AUF1, KSRP *etc*. are destabilizing proteins [27]. HuR is a ubiquitously expressed member of ELAV/Hu family proteins that are predominantly localized in the nucleus of the cell and under stressed conditions can shuttle from nucleus to the cytoplasm. Generally, the cytosolic HuR is involved in post-transcriptional and translational regulation of the target mRNAs [28, 29]. Numerous reports demonstrated that *beclin1* mRNA was downregulated by influence of miRNAs, like miR-30a, miR-17 and miR-20a [30–32]. miRNA mediated inhibition of Beclin-1 suppresses autophagy which can cause pathological conditions like cancer. On the other hand, inhibiting these miRNAs can restore autophagy.

Thus, the major aim of this study is to delineate the molecular mechanism by which expression of Beclin-1, one of the important players in WA-mediated impaired autophagy is regulated during WA-mediated impaired autophagy in MCF-7 cell-line.

## Results

### Post-transcriptional regulation of Beclin1 expression during WA-treatment in MCF-7 cells

To unravel the mechanism behind the impaired autophagy in breast cancer cell-line MCF-7 induced by WA treatment [13], the ratio of LC3B-II/ LC3B-I (a hallmark of autophagy) was measured by immunoblot (figure 1A). The result revealed that initially (3 h) LC3B-II level was increased upon WA-treatment (4 µM), but, 8 h onward the level of both LC3B-II and LC3B-I got increased as compared to control (24 h DMSO-treated). The expression of Becn1 at the level of mRNA (figure 1B) was found to be increased slightly (∼2 fold) at initial 3 h and remained almost same upto 24 h. However, the Becn1 protein level (figure 1C) was increased initially and it started to decline from 8 h onwards. It was found that 24 h after WA-treatment the level of Becn1 protein was ∼1.8 fold less as compared to DMSO-treated control. These results elicited that WA-treatment induced autophagy in the early hours which became impaired in the later hours similar to that reported by Ghosh et al. 2017 [13].

**Figure 1.**
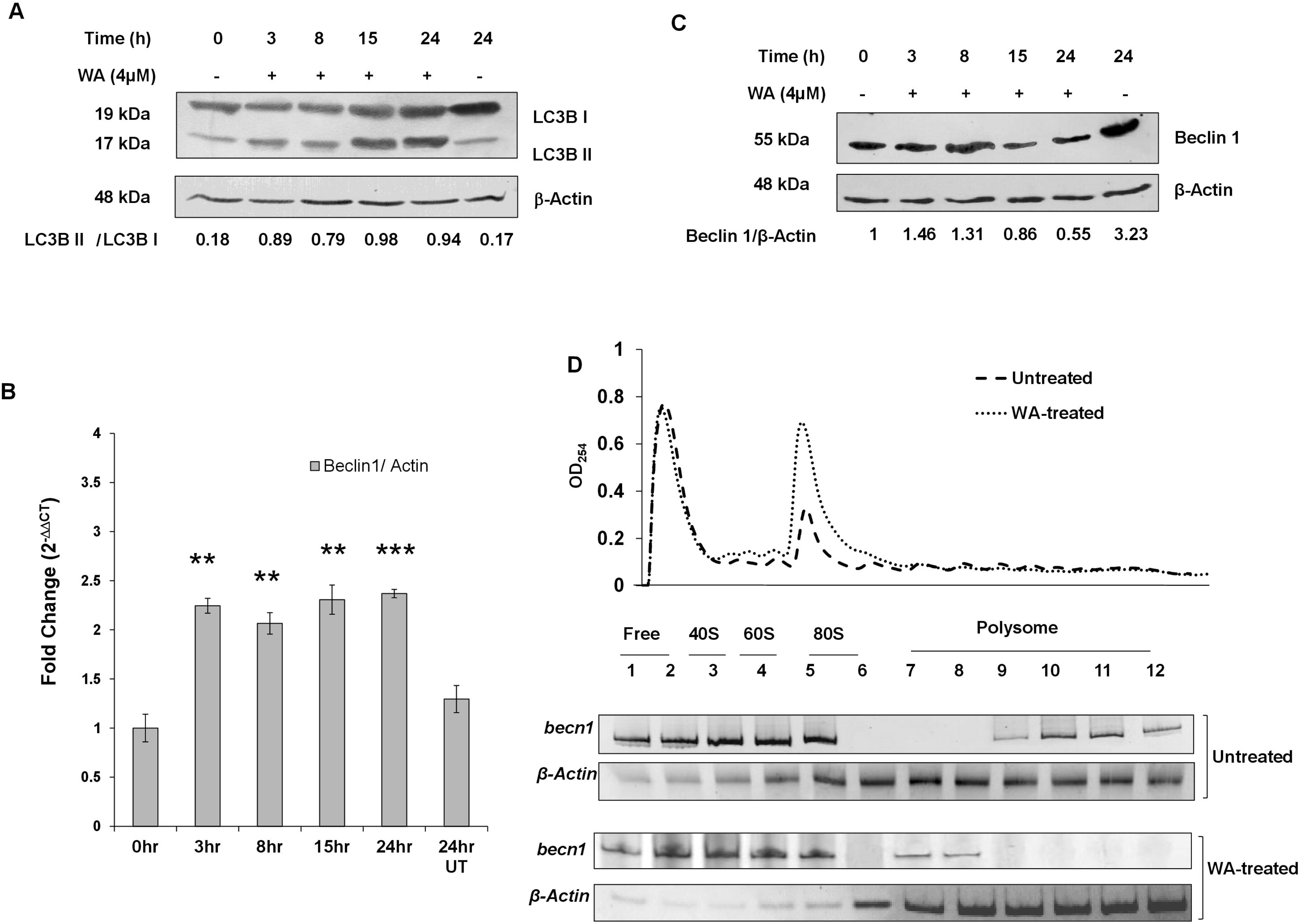
Post-transcriptional regulation of Beclin1 expression during WA-treatment induced impaired autophagy. (A) Western blot using total cellular proteins (80µg each) from WA-treated (4 μM for 24 h) MCF-7 cells, resolved in 14% SDS-PAGE and probed with LC3B specific antibody and anti β-Actin antibody (loading control). (B) Bar diagram showing relative expression (RNA level) of beclin1 mRNA level during WA-treatment induced autophagy at indicated time points measured by qRT-PCR using ΔΔC_T_ method and normalized with β-actin mRNA. This plot for quantitative PCR is mean of three independent experiments and presented as mean□±□SEM where NS indicates (P□>□.05), where * is (P□≤□.05), ** is (P□≤□.01) and *** is (P□≤□.001). (C) Western blot showing Beclin1 protein level during WA-induced autophagy where cellular proteins (30 µg) were resolved in 12% SDS-PAGE and blotted with Beclin1 specific primary antibody and re-probed with anti β-Actin antibody (loading control). (D) Polysome profiles of untreated and WA treated MCF-7 cells (top). Semi-quantitative RT-PCR of becn1 mRNA and β-actin from polysome fractions of untreated and WA treated MCF-7 cells (bottom). Polysome profiling was repeated twice.

Translational yield (TY) was found to be reduced (0.2) due to WA-treatment which claimed the possibility of repression in *beclin1* protein synthesis (Fig. S1). To probe the reason, polysome profiling was performed (figure 1D; upper panel). mRNA levels of *becn1* and β*-actin* were measured by semi-quantitative RT-PCR (figure 1D; lower panel) from fractions obtained after sucrose density gradient centrifugation of DMSO-treated and WA-treated cytosolic extracts of MCF-7 cells. The results indicated that *becn1* mRNAs were less associated with the polysome in WA-treated cells as compared to the DMSO-treated ones.

### Involvement of microRNA-20a in the expression of Beclin1

*In silico* search revealed a few highly conserved regions in the 3’-UTR of *becn1* mRNA. The entire 628 bases long *becn1* 3’-UTR was scanned for putative miRNA target sites using public web-based miRNA target prediction software: Target Scan Human version 7.0, which revealed the presence of seed regions for several families of miRNAs.To check the *in vivo* association of any microRNA (miRNA) to 3’-UTR of *becn1* mRNA, co-immunoprecipitation was performed with anti-Ago2 antibody using DMSO- or WA-treated (24h) cell extracts. Figure 2A-i; upper panel has displayed a band of 200 bp PCR product specific for *becn1* mRNA in both the inputs and WA-treated IP sample, where no such band was detected in DMSO-treated IP sample. IPs with normal IgG antibody (negative control) for both the conditions showed no such bands, where amplification of β*-actin* RNA was observed only in the inputs indicating specificity of IP. Western blot with the same samples (figure 2A-i; lower panel) confirmed the success of the immunoprecipitation process. The above-mentioned results clearly demonstrated that Ago2 was associated with *becn1* mRNA only in WA-treated MCF-7 cells but not in control cells. The result was also confirmed quantitatively by RT-qPCR analysis of the same samples (figure 2A-ii). However, no change in the Ago2 protein expression by WA-treatment was established by immunofluorescence (figure S2).

**Figure 2.**
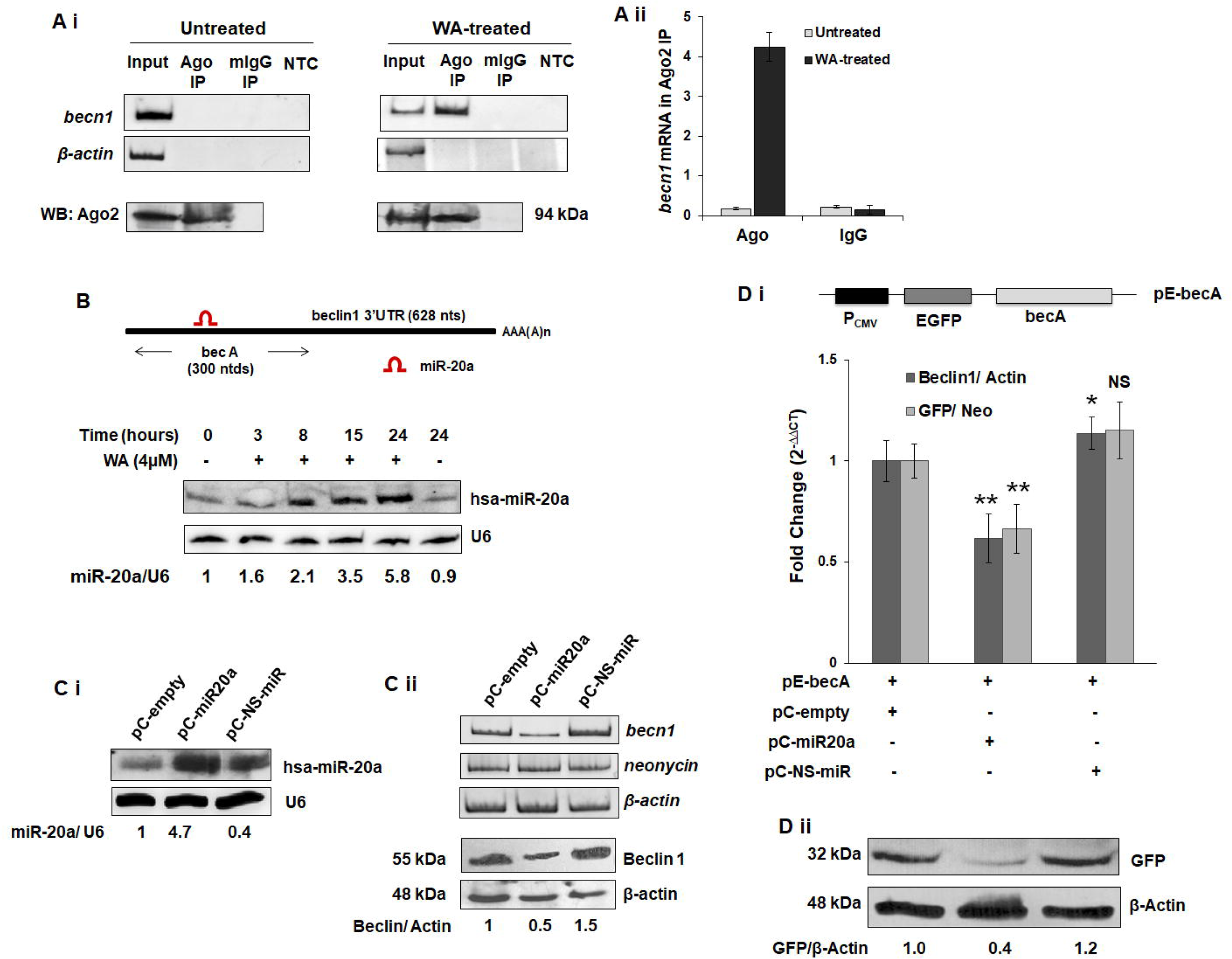
microRNA-20a is engaged in post-transcriptional regulation of Beclin1 expression. (Ai) Elucidation of interactions of *bec-A* mRNA with Ago2 *in-vivo*. Upper panel: Ethidium bromide-stained gel picture of semi-quantitative RT-PCR products of *bec-A* and β*-actin* RNA of chemically crosslinked RNA-protein complexes from untreated and WA treated MCF-7 cells immunoprecipitated with anti-Ago2 or IgG antibodies. Lower Panel: Western blot with anti-Ago2 of immunoprecipitated samples (as indicated). [IP-immunoprecipitate; NTC-No template control]. (A ii) Quantitative analysis of becn1 mRNA immune-precipitation with Ago2 or IgG antibodies of untreated or WA treated MCF-7 cell lysates. The relative expression was calculated by ΔΔC_T_ method and normalized with β-actin mRNA. Plotted data are means of three independent experiments and presented as mean□±□SEM. (B) Upper panel is illustrating the schematic representation of beclin1 3 UTR (not in scale) emphasizing the target hybridization site of microRNA-20a or miR-20a on proximal part of *becn1* 3’-UTR or *bec-A*. Lower panel is showing the northern blot analysis of mature-miRNAs in WA-treated (4 μM for 24 h) MCF-7 cells at indicated time points. 15μg of total RNAs were resolved in 15% 7(M) Urea-PAGE andmature microRNA levels were analyzed by blotting with radioactive [^32^P] probes. The same blot was reprobed with U6 (loading control). (C i) Northern blot with U6 probe used as a loading control. (C ii) Upper panel shows intrinsic beclin1 mRNA level in miR-20a over-expressed MCF-7 cells, measured by semi-quantitative PCR after staining with ethidium bromide. *Neomycin* was used as transfection control and β-actin as loading control. Lower panel: Beclin1 (intrinsic) protein level with β-Actin level as loading control of miR-20a over-expressed MCF-7 cells. (D i) MCF-7 cells co-transfected with pE-Bec-A and pc-empty/ pC-miR20a/ pC-NS-miR constructs for 48 h. Relative expression of beclin-1 or gfp reporter RNA was measured by qRT-PCR (ΔΔC_T_), normalized to β-actin and neomycin mRNA respectively. (D ii) 30μg of proteins extract from miR-20a over-expressed MCF-7 cells were resolved in 12% SDS-PAGE and were analyzed by western blot using anti-Beclin1 antibody, anti-β-Actin antibody (loading control). Data plotted in A ii and D i are means of three independent experiments and presented as mean□±□SEM where NS indicates (P□>□.05), * is (P□≤□.05), ** is (P□≤□.01) and *** is (P□≤□.001).

As the seed sequence for hsa-miR-20a (135-141 nucleotide downstream of stop site) (figure 2B; upper panel) is present in the *becn1* 3’-UTR, the expression level of mature miR-20a was determined by northern blot (figure 2B; lower panel). It was found that the level of has-miR20a was increased steadily in MCF-7 cells with increasing time of WA treatment. Ratio of the expression of miR-20a *w.r.t.*U6 (loading control) was found to be increased by 5.8-fold.

To mimic the WA-treated condition (*i.e.* increased expression of miR-20a), pC-miR-20a (sense strand of pre-miR-20a [33]) was overexpressed in MCF-7 cells by transfection, where, cells transfected with empty-vector (pC-empty) and unrelated miRNA (pC-NS-miR) were used as transfection control and negative control respectively. The mature hsa-miR-20a expression levels for all three sets were checked by northern blot (figure 2C-i), illustrating higher level of hsa-miR-20a expression in the cells transfected with pC-miR-20a as compared to the other two controls, where U6 level for each sample indicting equal load of RNA in each lane. To find out the effect of miR-20a on beclin1 expression, intrinsic *becn1* mRNA was measured by RT-PCR. Figure 2C-ii; upper panel showing decreased level of intrinsic *becn1* mRNA in pC-miR-20a transfected cells as compared to both the controls (pC-empty and pC-NS-miR). Level of *neomycin* mRNA and β*-actin* indicating efficiency of transfection and amount of loading respectively. Figure 2C-ii; lower panel elucidated reduction of endogenous level of Becn1 protein (as measured by western blot with anti-Becn1 antibody) by over-expression of miR-20a in MCF-7 cells.

To ascertain the direct interaction of miR20a with the 3’-UTR of *becn1* mRNA proximal region of *becn1* 3’-UTR (300 bp) containing miR-20a seed sequence (named as *bec-A*, schematically showed in figure 2B; upper panel) was cloned in pEGFPC1 at the 3’-end of *gfp* gene to yield pE-bec-A (figure 2D-i; top). MCF-7 cells were co-transfected with pE-bec-A and pC-empty/pC-miR-20a/pC-NS-miR vectors. Total RNAs and total proteins were extracted from those co-transfected MCF-7 cells and the level of *gfp* mRNA and protein was analyzed by RT-qPCR and western blot respectively. The results showed that the *gfp* mRNA level was decreased by ∼1.6 fold and GFP protein level by 2.5-fold in pC-miR-20a overexpressed cells as compared to pC-empty transfected cells (figure 2D). The above result indicated that miR-20a could modulate the expression *becn1* mRNA by associating with the *bec-A* region at its 3’-UTR.

### Association of HuR with the 3’-UTR of beclin1 mRNA

*In-silico* characterization of *becA* predicted putative *cis*-elements for binding of RNA binding proteins (RNA-BPs), *e.g.* Class I (AUUUA) and Class III (UUUUAAAAUUAA) AU-rich Elements (AREs) positioned close to seed sequence of miR-20a family members (figure 3A). Several reports have established that AREs are the binding sites for various RNA stabilizing (Nucleolin, HuR*etc.*) and de-stabilizing (TTP, AUF1*etc.*) proteins [18]. For identification of *bec-A* interacting protein(s) oligo dT affinity chromatography was performed using *in vitro* transcribed, poly-adenylated *bec-A* transcript and WA-treated cytoplasmic extracts of MCF-7 cells. Figure 3B-i is showing coomassie blue stained SDS-PAGE profile of proteins present in the eluted fractions from *bec-A* affinity chromatography. Notably, a major protein band of molecular weight ∼36 kDa was apparent in fractions eluted with 200 to 800 mM of NaCl (figure 3B-i). Western blot (figure 3B-ii) revealed the presence of a known RNA binding protein, HuR (36 kDa) in the eluted fractions (eluted with 400and 600 mM NaCl) however, the specificity of this interaction was confirmed by the absence of β-Actin in the eluted fractions (while being present in the input).

**Figure 3.**
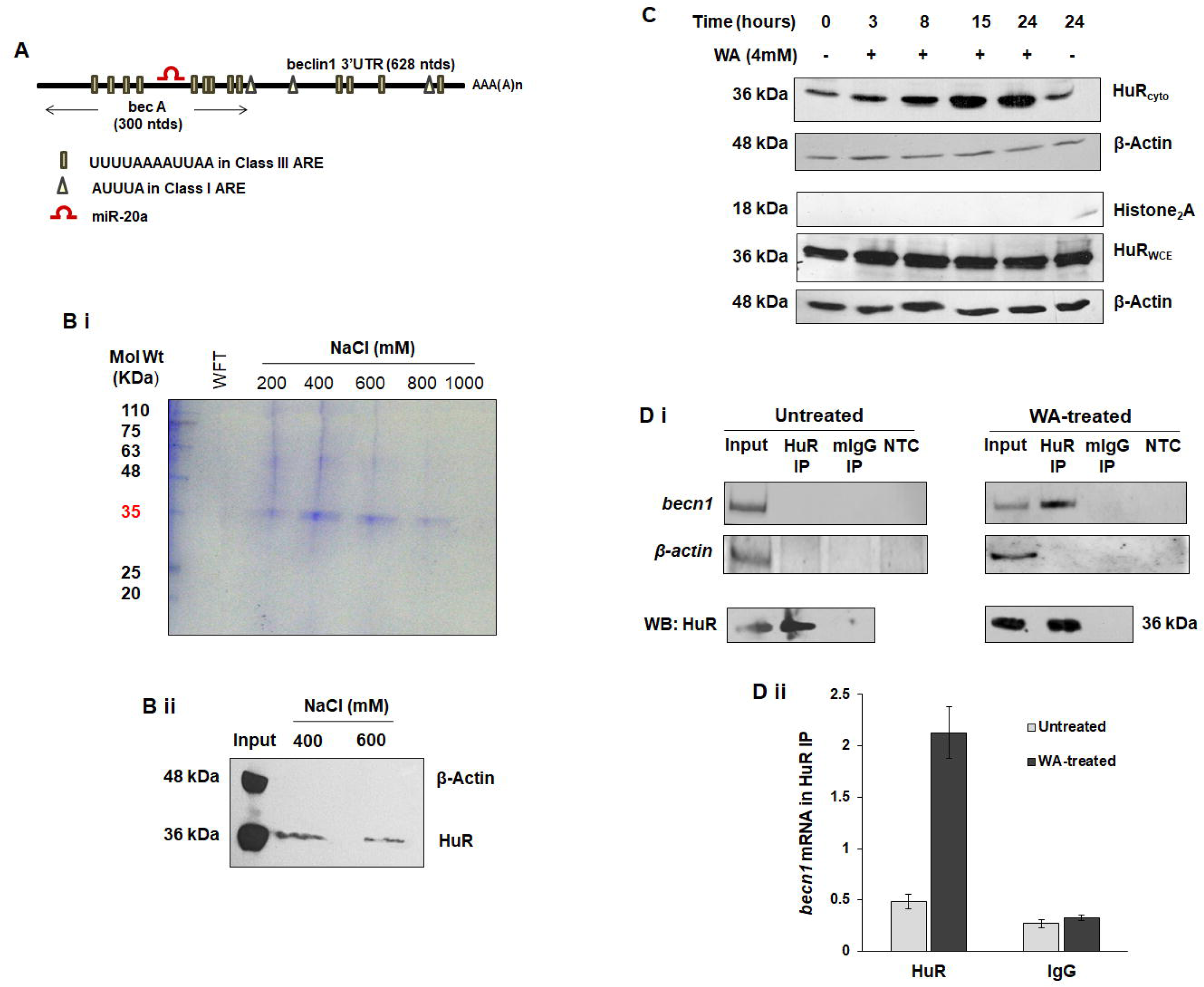
Interactions of HuR with proximal part of beclin1 3’-UTR. (A) In-silico characterization of becn1 3’-UTR (not in scale). Schematic representation shows the proximal part of *bec-A* on 3’-UTR based on the positions of AU rich elements and microRNA-20a seed sequences. (B i) Elucidation of interactions of *bec-A* mRNA with HuR *in-vitro*. Elution profile of the proteins in WA-treated MCF-7 cytoplasmic extracts eluted from the RNA affinity (polyadenylated-*bec-A* bound oligo-dT beads) column, where eluted fractions were separated in 10% SDS-PAGE and stained with Coomassie Blue. (B ii) Western blot of column fractions eluted with NaCl and blotted with anti- HuR and anti-β-Actin antibodies. [Abbreviations: MW- molecular weightmarkers; LFT-Loading flow through; WFT-wash flow through]. (C) Western blots of HuR with MCF-7 cells untreated or WA-treatment, with whole cell and cytoplasmic extracts where β-Actin was used as loading control and Histone-2A to show no nuclear contamination in cytosolic extracts. 30μg of total proteins were resolved in 12% SDS-PAGE. [Abbreviations: CE-cyto-cytoplasmic extracts; WCE-whole cell extracts]. (D i) Elucidation of interactions of *bec-A* mRNA with HuR *in-vivo*. Upper panel: PCR products obtained by semi-quantitative RT-PCR using bec-A and β-actin specific primers from chemically crosslinked RNA-protein complex IP with HuR or IgG antibodies from untreated, or WA-treated MCF-7 cells separated on 6% native PAGE. Lower Panel: western blot of immunoprecipitated samples (as indicated) with anti-HuR. [IP: immunoprecipitate; NTC: No template control]. (D ii) Quantitative analysis of becn1 mRNA immune-precipitation with Ago2 or IgG antibodies of untreated or WA treated MCF-7 cell lysates. The relative expression was calculated by ΔΔC_T_ method and normalized with β-actin mRNA. Plotted data are means of three independent experiments and presented as mean□±□SEM where NS indicates (P□>□.05), * is (P□≤□.05), ** is (P□≤□.01) and *** is (P□≤□.001).

HuR was reported as predominant nuclear protein [29], to claim its interaction with the cytosolic mRNAs, presence of HuR in the cytosol is required. Western blot (figure 3C) has revealed WA-driven time-dependent increase of cytosolic HuR, with no considerable change in its total cellular level, suggesting a nuclear to cytosolic translocation of HuR. The presence of an insignificant amount of histone indicated absence of nuclear protein contamination in the cytosolic extract.

To confirm the *in vivo* association of HuR to *becn1* mRNA in WA-treated MCF-7 cell, co-immunoprecipitation (co-IP) with specific antibody against HuR was performed. Presence of *becn1* mRNA in HuR-IP sample were estimated by semi-quantitative RT-PCR, where the presence of a band of 200 bp amplicon in HuR-IP sample of WA-treated MCF-7 cells and its absence in untreated one (Figure 3D-I; upper panel) confirmed the specific interaction of HuR with *becn1* mRNA in the WA-treated cells. Mouse-IgG antibody was used as negative control for IP while presence of β*-actin* band in input and its absence in IP samples indicated binding specificity of HuR to *becn1* mRNA. Success of IP was confirmed by western blot with HuR antibody (figure 3D-i; lower panel). Similar observation was found by quantitative RT-PCR analysis of the same samples (figure 3D-ii).

### Involvement of GW-body in Beclin1 Translational inhibition

GW-182 (glycine-tryptophan protein of 182 kDa) is an essential component of GW-body subunit, participates in miRNA-mediated gene silencing [34] and it helps to localize the mRNAs in P-/GW-body [35]. Semi-quantitative (figure 4A-i) and quantitative (figure 4A-ii) RT-PCR revealed that *becn1* mRNA was co-immunoprecipitated with GW-182 antibody in WA-treated condition only.

**Figure 4.**
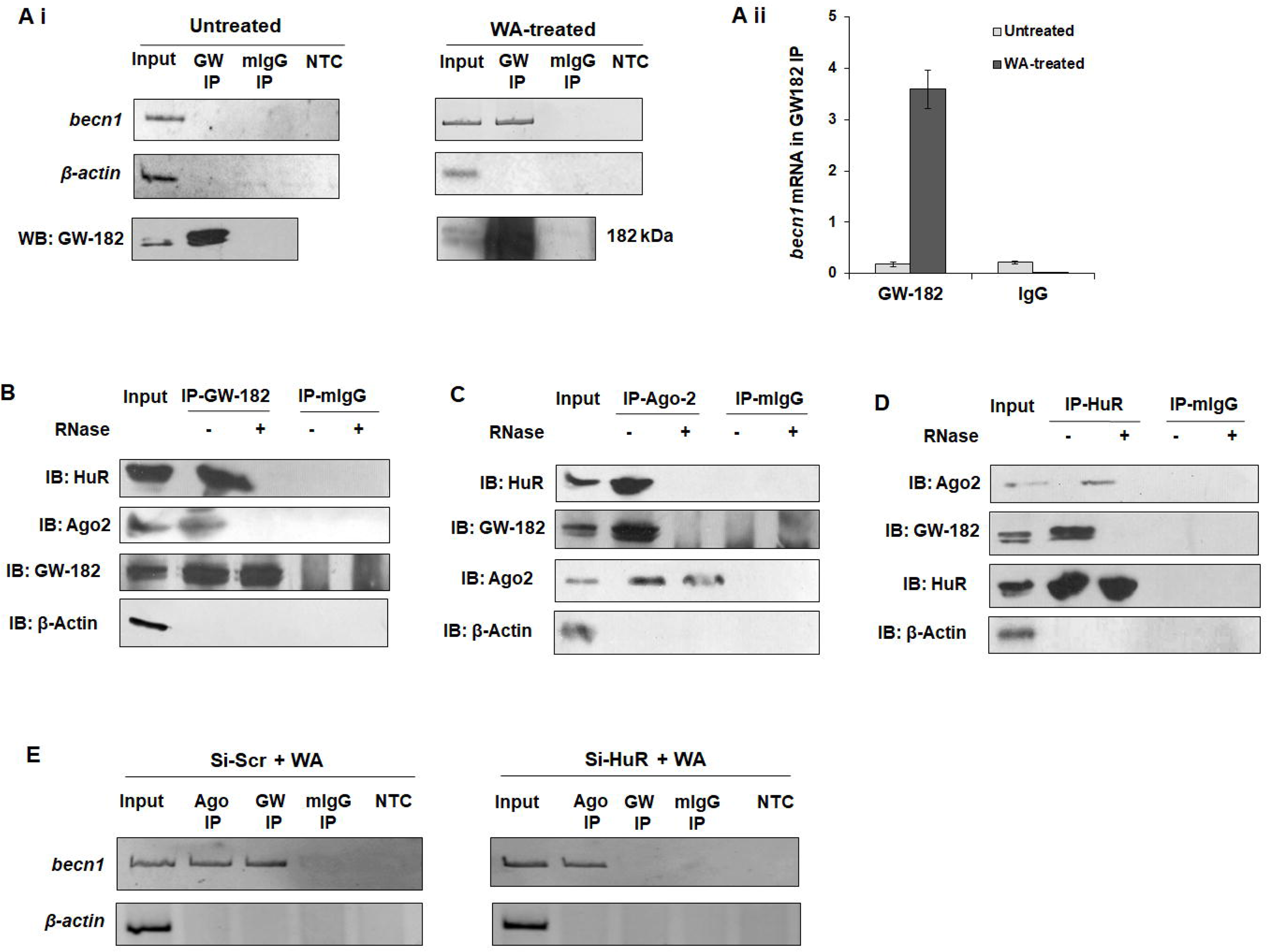
Beclin1 translational inhibition continues with GW-body association. (Ai) Elucidation of interactions of *bec-A* mRNA with GW-182 *in-vivo*. Upper panel: Ethidium bromide stained gel picture of semi-quantitative RT-PCR products of bec-A and β-actin RNA of chemically crosslinked RNA-protein complexes from untreated and WA treated MCF-7 cells immunoprecipitated with anti-GW-182 or IgG antibodies. Lower Panel: Western blot with anti-GW-182 of immunoprecipitated samples (as indicated). (A ii) Quantitative analysis of becn1 mRNA immune-precipitation with GW-182 or IgG antibodies of untreated or WA treated MCF-7 cell lysates. The relative expression was calculated by ΔΔC_T_ method and normalized with β-actin mRNA. Plotted data are means of three independent experiments and presented as mean□±□SEM where NS indicates (P□>□.05), * is (P□≤□.05), ** is (P□≤□.01) and *** is (P□≤□.001). (B, C and D) Western blots for proteins from IP with GW-182, Ago2, HuR or IgG antibodies of WA-treated MCF-7 cytoplasmic extracts pre-incubated with or without RNase A. Proteins were resolved in10% SDS-PAGE. (E) Elucidation of interactions of *bec-A* mRNA with Ago2 and GW-182 *in-vivo* after depletion of HuR. Ethidium bromide-stained gel picture of semi-quantitative RT-PCR products of *bec-A* and β*-actin* RNA of chemically crosslinked RNA-protein complexes from untreated and WA treated MCF-7 cells with scramble siRNA (si-Scr) or HuR siRNA (si-HuR) immunoprecipitated with anti-Ago2, anti-GW-182 or IgG antibodies. [Abbreviations: IP-immunoprecipitate; NTC- No template control].

In order to reveal whether HuR protein and any microRNA is associated with GW-182 protein, protein-protein co-immunoprecipitation assay (figure 4B) was performed using anti-GW-182 antibody to pull down the complex from WA-treated MCF-7 cell extract followed by immuno-blotting with anti-HuR and anti-Ago2 antibodies. It was observed that GW-182 interacts with HuR and Ago2 in WA-treated condition. Similarly, the presence of HuR protein and GW-182 in the complex pulled down with anti-Ago2 antibodies (figure 4C) and the presence of GW-182and Ago2 in the complex pulled down with anti-HuR antibodies (figure 4D) pointed towards the fact that in WA-treated MCF-7 cells HuR, GW-182 and Ago2 (microRNA) stays associated with each other. This association was however perturbed when the cytoplasmic extract was treated with RNase A/T1 prior to pull downconfirming an RNA-dependent interaction between these three proteins in cytoplasm of WA-treated MCF-7 cells (figure 4A, 4B, 4C).

Knocking down of HuR by gene-specific siRNA (si-HuR) followed by co-IP (figure 4E) revealed that 200 bp band of *becn1* was present in IP sample with anti-Ago2 but absent in anti-GW-182 sample in WA treated MCF-7 cells. In contrast, using non-specific siRNA (si-Scr) showed presence of the band in both cases. The results revealed that silencing of HuR released GW182 from *beclin1* 3’UTR, while Ago2 interaction remains in WA-treated cells. This implies the role of HuR in transport of *beclin1* mRNA entangled with miR-20a/Ago2 to GW bodies.

## Discussion

Autophagy is a vital cellular homeostasis process. Deregulated expression of the autophagy related genes or viral infections or anti-cancer agents can block autophagic pathways at different levels facilitating impairment in autophagy that often can lead to diseases [7, 10, 36]. Withaferin-A (WA) was reported to have autophagy inducing proficiency [37]. Our group has established that WA caused impaired or defective autophagy in breast cancer cell-lines due to reduction in the level of upstream autophagy markers, like BECN1, one of the essential genes for autophagosome formation [13]. Thus, exploring the mechanism of regulation of BECN1 expression in incomplete autophagy became the key goal of this study.

In the present study, it was observed that the BECN1 protein level got reduced with increasing time of WA treatment, though the *becn1* mRNA level remained almost unchanged. The calculated translational yield (TY) of BECN1 in WA-treated MCF-7 cells was found to be reduced from 1 to 0.2 (after 24 h). It was also found that the association of *becn1* mRNA with the polysomes was also got reduced indicating a possibility of inhibition in protein synthesis (figure S1). miRNAs are well-studied key negative regulator of gene expression that can promote mRNA degradation and inhibition of translation as well [38]. Ago2, one of the key components of RISC (RNA-induced silencing complex) plays crucial role in recognizing the specific sequences in the 3’-UTR of its target mRNAs and recruits the miRNAs [39]. Generally, the RNA-IP with Ago2 followed by microarray analysis confirms the association of active pool of miRNAs with their corresponding targets (miRNA targetomes) [40]. Here, RNA co-IP using anti-Ago2 followed by semi-quantitative RT-PCR established more association of miRNA(s) with *becn1* mRNA in WA-treated MCF-7cells as compared to untreated control.

To identify the specific miRNAs involved in this process, *in-silico* analysis was used to foresee the probable binding sites for microRNAs on *becn1* 3’-UTR, where hsa-miR-20a was predicted with the highest context score. Interestingly, in our case, it was found that the mature miR-20a level was high and the endogenous Becn1 protein level was low in WA treated MCF-7 cells as compared to the control cells. It was also evident from the results that miR-20a over-expression (which mimic the WA-treated condition) has negative effect on *becn1* expression. The reason for the apparent discrepancy in this result (figure 2Cii: upper panel) with figure 1B (slight increase of becn1 mRNA with WA treatment) could be that, WA-treatment caused translocation of HuR to the cytoplasm, which could bind to the *becn1* mRNA thus preventing the decrease in its level. Additionally, *gfp*-reporter assay elucidated that miR-20a exclusively interacted with the seed sequence present in the 3’-UTR of *becn1* mRNA resulted in downregulated BECN1 expression. There might have some contribution of miRNAs in the translational repression of *becn1* in WA-induced impaired autophagy. Thus, this study may also shed light on the specific mechanisms of target silencing by miRNAs under non-canonical form of autophagy in general.

Other than miRNAs, RNA Binding Proteins (RBPs) that interact with mRNAs are also major players in determining the fate of an mRNA. High-throughput screen identified the eukaryotic translation initiation factor, eIF5A as a positive regulator of autophagy that promote translation of *ATG3* [41]. Another translation initiation factor, eIF4E-BP1 was found to act as negative regulator of autophagy [42] in maintaining the proper homeostasis of cells. HuR, the most studied member in ELAV protein family, is largely reported for its role in mRNA stability and translation, although it participates in transcription, alternative splicing, polyadenylation and subcellular localization of target mRNAs [43, 44]. Recently, our group had established that HuR promotes stabilization and translation of *becn1* during serum starvation induced autophagy in MCF-7 cells [28]. Other than *becn1*, HuR also binds to 3’-UTR of *ATG5*, *ATG12*, and *ATG16* mRNAs and mediates their translation in HCC cells to promote autophagosome formation [45]. In pathology of age-related macular degeneration, HuR was found to suppress autophagic degradation by promoting SQSTM1/p62 synthesis [46–47]. In the current study, RNA affinity chromatography and co-IP using HuR antibody confirmed association of mRNA stabilizing factor, HuR on *becn1* 3’ UTR in WA-treated MCF-7 cells. Although HuR typically stabilizes and/or increases target mRNA translation, there are handful examples indicating repressive effect of HuR on target mRNAs [48–50].

Simultaneous interaction between Ago protein bound miRNAs and RNA Binding Proteins (RBPs) with target mRNAs determine the outcome of gene regulation in both physiological conditions and in pathology [51]. This kind of combinatory regulation is observed when HuR and Ago2-miR complex interact with the mRNAs and functions either in competitive or cooperative manner [52–53]. Here we have demonstrated the simultaneous interaction of miR-20a/Ago2 and HuR with 3’-UTR of *becn1* mRNA and thus could be responsible for reduced expression of Beclin1 protein in WA-treated MCF-7 cells. Hyeon et al., 2009 established that HuR facilitates the interaction of let-7-RISC with the 3’UTR of *myc*, leading to its destabilization and translational repression [54]. In contrary, HuR deficiency increases the level of miR-466i expression in Th17 cells to mediate GM-CSF and IL-17 mRNA decays [55].

Several reports showed that the association of miRNAs/AGOs and HuR might result in opposing effects on the localization of mRNAs, either in translationally active polysomes or in repressive cytoplasmic complexes e.g., Processing-Bodies or Stress-Granules [56–58]. For example, HuR binding to 3’-UTR of CAT-1 mRNA relocates it from PBs (P-bodies or Processing bodies) to cytoplasm for re-association with polysomes in Huh7 cells, which helps to overcome the miR-122/AGO mediated translational inhibition [56]. More reports clarified that HuR promotes the translation of occludin and E-cadherin mRNAs by releasing them from PBs [59–60]. On the other hand, HuR stabilizes target mRNAs and transfers them in SGs or stress granules in response to environmental stress [61]. Likewise, WA-treatment promotes localization of HuR associated miR-20a entangled *becn1* mRNA to GW-182, whereas, knocking down of HuR reduces the affinity of GW-182 protein towards *becn1* 3’-UTR. In the presence of RNase A/T1 this association was however entirely disconcerted.

Nevertheless, the studies on RBP mediated post-transcriptional and translational regulation of ATG genes are comparatively less explored. Here the observed interactions of the 3’-UTR of *becn1* mRNA in WA-treated MCF-7 cells with RBPs are schematically summarized in figure 6. Co-immunoprecipitation using Ago2 antibody established *in vivo* association of miRNAs with *becn1* mRNA, whereas HuR, a predominant nuclear protein that after translocating to the cytosol and co-IP using GW-182 antibody confirmed the interaction of GW-182, a major component of cytosolic GW-body associate with *becn1* 3’ UTR in WA-treated MCF7 cells simultaneously (figure 5 I, II, III). From this study it can be concluded that the fate of *becn1* mRNA had been decided by simultaneous binding of miR-20a (miRNA) Ago2 and HuR (RBP). Under WA-treated condition, binding of both *trans*-factors at a time stabilizes *becn1* mRNA and transfer them to stress granules (SGs), thus prevent the association of this mRNA to the translational machinery and downregulate the expression of Beclin1 protein. The scarcity of Beclin1 protein hindered autophagosome formation and triggered WA-mediated impaired autophagy in MCF-7 cells. This conclusion has been pictorially represented in the Figure 6.

**Figure 5.**
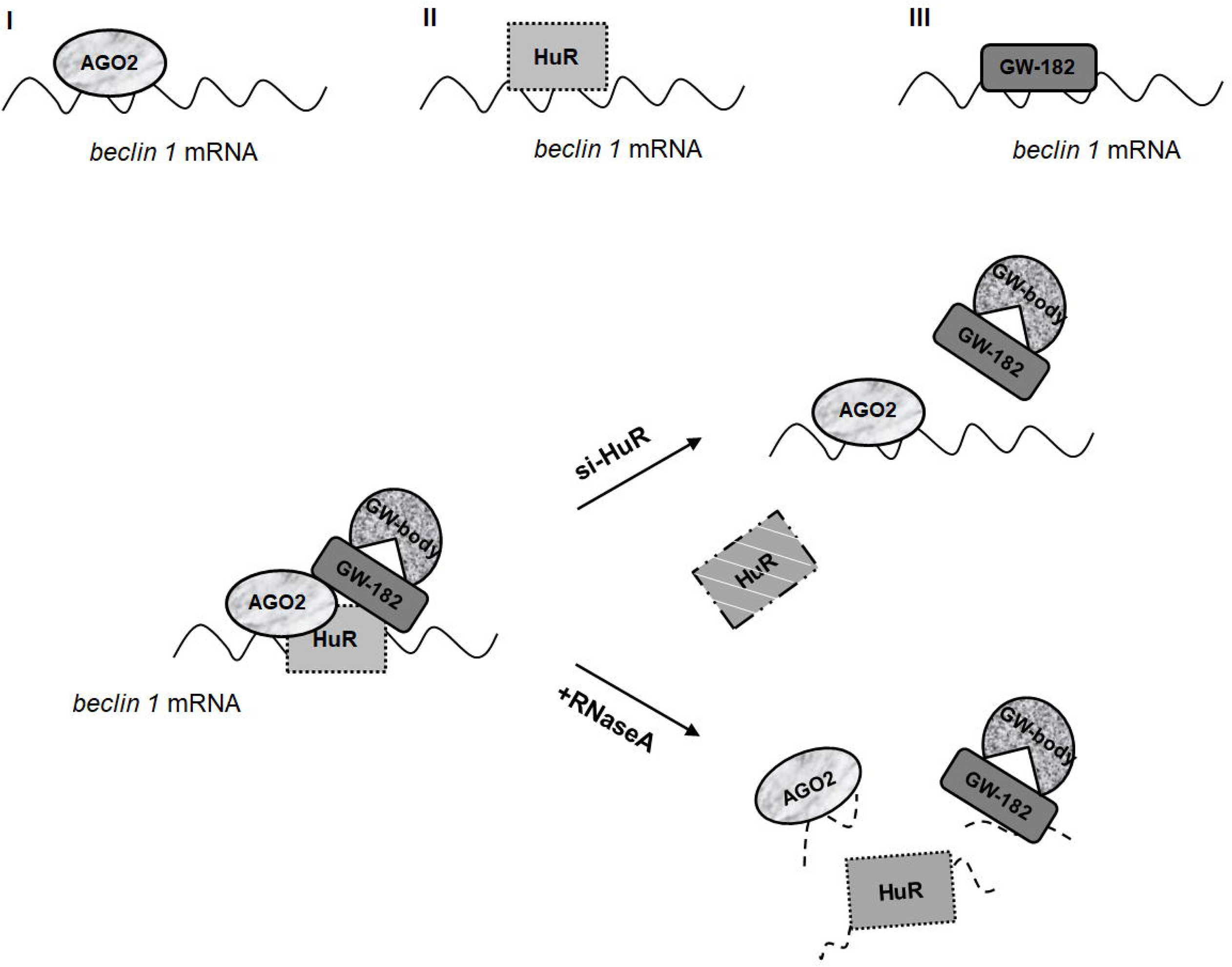
Schematic diagram showing interaction between beclin1 mRNA with RBPs. Interactions of the 3’-UTR of *becn1* mRNA in WA-treated MCF-7 cells with Ago2, HuR and GW-182 has been schematically summarized.

**Figure 6.**
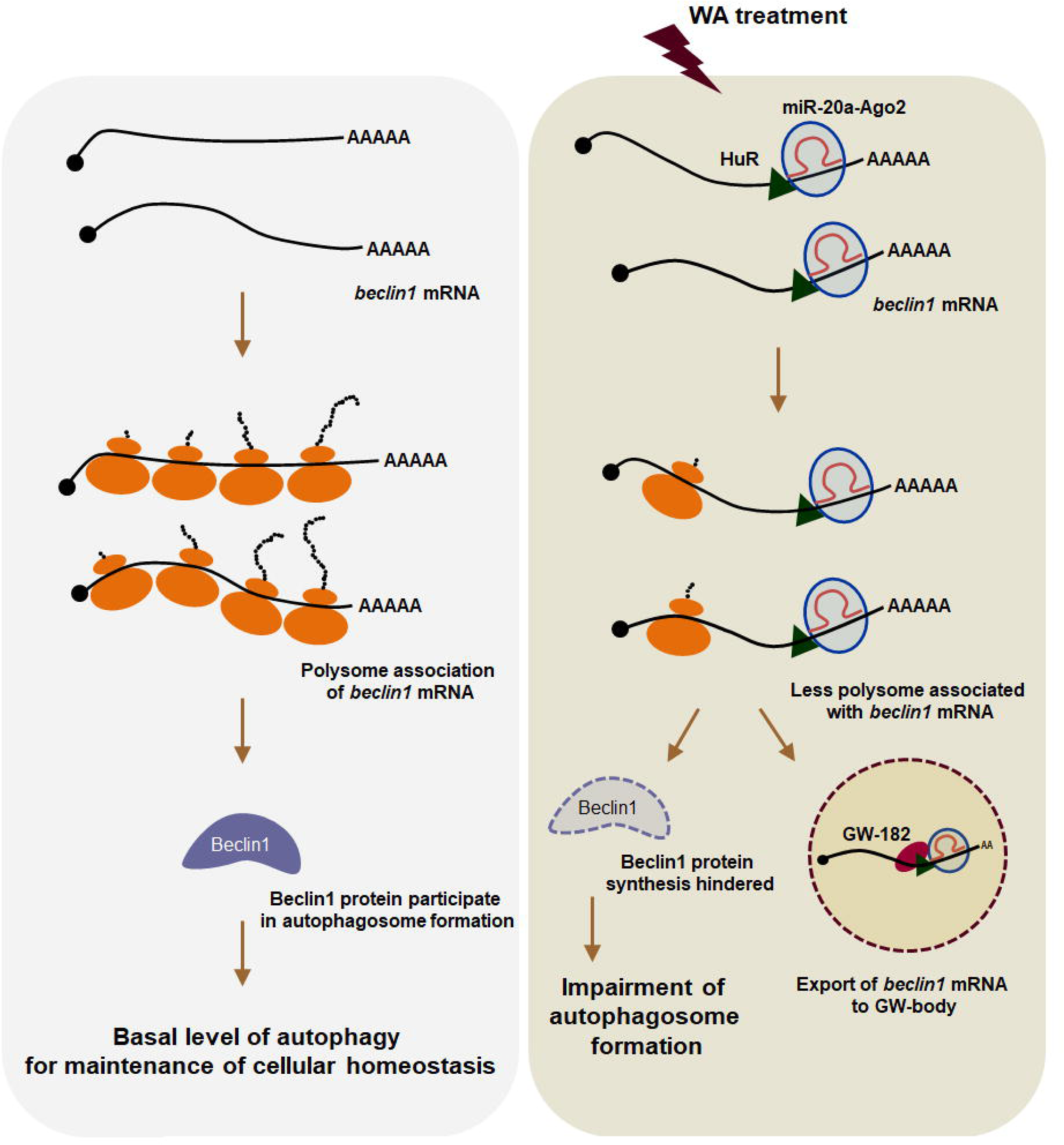
Schematic diagram showing regulation of cellular autophagy in WA-treated MCF-7 cells. Modulation of Beclin1 through inhibition of translation by sequestration of its mRNA with the help of Ago2 bound miRNA-20a, HuR and GW-182 proteins disrupt autophagy in WA-treated MCF-7 cells.

## Materials and Methods

### Cell culture and treatment with Withaferin-A

Human epithelial breast carcinoma cell-line, MCF-7 was obtained from National Centre for Cell Science, Pune, India and maintained as described in [13]. MCF-7 cells (0.8-1X10^6^) were treated with 4μM WA (Alexis Biochemicals, USA) for 0-24 h in 37°C and harvested at various time points to obtain cellular RNA or protein. DMSO was used as solvent to dissolve WA.

### In silico study

3’UTR of *becn1* (628 nucleotides long) was scanned for putative miRNA target sites using public web-based miRNA target prediction software namely miRanda-miR SVR (microRNA.org) and Target Scan Human 7.2 (http://www.targetscan.org) that identified a binding site of mature hsa-miR-20a. The *in-silico* study also revealed the conservation of miR-20a seed sequence among the vertebrates (http://www.targetscan.org). Proximal region of *becn1* 3□-UTR containing miR-20a seed sequences was mentioned as *bec-A* (300 nucleotides in length).This region of 3’-UTR or *bec-A* was also characterized *in silico* with respect to their A+U contents.

### Cloning

A region of 3’-UTR of *becn1* mRNA [nucleotides# 13534-13834, (NM_003766.3)] was cloned in pTZ57R/T (Thermo Scientific, USA) by RT-PCR using primer pair (5’-CTT-TTT-TCC-TTA-GGG-GGA-G-3’/ 5’-CAA-CTC-AGT-TAA-AAA-AAA-GAA-AAG-C-3’) to produce pT-bec-A. The clone was confirmed by sequencing in Gene Analyzer 3130 (Thermo Scientific, USA). This region was further digested with specific restriction enzymes and subcloned into the mammalian expression vector pEGFPC1 downstream *gfp* reporter gene to produce pE-bec-A. The cloning of pC-miR-20a and pC-NS-miR are mentioned in [62].

### Transient Transfection of plasmids and siRNAs

MCF-7 cells (3X10^5^) were seeded with antibiotic free fresh media in 35 mm tissue culture plates. pC-miR-20a (in pC-DNA 3.1 vector), pC-HuR and pE-becA constructs were transiently transfected using Turbofect transfection reagent (Thermo Scientific) according to manufacturer’s protocol.

For siRNA transfections, 50nM of control siRNA or HuR sense 5□-CCA-GUU-UCA-AUG-GUC-AUA-A55-3□ and anti-sense 5□-UUA-UGA-CCA-UUG-AAA-CUG-G55-3□ duplex RNA (Eurogentec, Belgium) were transfected using jetPRIME plasmid/siRNA transfection reagent (Polyplus-transfections, Illkirch, France) according to manufacturer’s protocol. 48h after transfection, the cells were harvested for isolation of total RNA or protein.

### Detection of autophagic vacuoles in MCF-7 cells stably transfected with ptfLC3 vector by confocal microscope

MCF-7 cells were stably transfected with ptfLC3 vectors (Addgene, MA, USA) and single colony (expressing both *gfp* and *rfp*) was isolated by limiting dilution in 96-well plates and were maintained in individual flasks in presence of 50µg/ml of G418 [13].

MCF-ptfLC3 cells were transfected with either pCDNA empty vector or pC-miR-20a, pC-NS-miR prior to WA treatment. The cells were permeabilized with 0.1% Triton-X100 for 10 min, mounted with n-propyl galate and were visualized by Olympus confocal laser scanning microscope (Ix81) at 40X magnification after staining with DAPI (5 ng/ml).

### RNA Extraction and qRT-PCR

Semi-quantitative and quantitative RT-PCR [using KAPA SYBR FAST qPCR kit in Step One Plus System (Applied Biosystems)] were performed using *becn1* specific primer pair (5’-GAA-ATT-TCA-GAG-ATA-CCG-ACT-TGT-TC-3’/5’-CCT-TTC-TCA-ACC-TCT-TCT-TTG-AAC-3’) with the cDNAs obtained from total cellular RNAs of MCF-7 cells according to [63]. β*-actin* and *neomycin* mRNAs levels were estimated as internal and transfection control respectively. For the reporter assays, level of *gfp* RNA were quantified by qRT-PCR and the relative expression levels were analyzed by ΔΔC_T_ method. Primer sequences for neomycin, β*-actin* and *gfp* are mentioned in [29].

### Western Blot Analysis

Whole cell and cytoplasmic extracts of DMSO- and WA-treated MCF-7 cells were used to perform western blots using antibodies against Beclin1, LC3, HuR, β-actin (Santa Cruz) and GW-182, histone, Ago2 (Cell Signaling).

### Preparation of RNA transcripts and microRNA probes

*becA* transcript was synthesized from pT-bec-A plasmid templates using T7 RNA polymerase. ^32^P-labeled RNAs were synthesized in transcription reactions containing ^32^P-CTP. To maximize the amount of full-length product, reactions were performed in presence of 250μM unlabeled CTP, along with 500µM each of ATP, UTP and GTP. The purity of [^32^P]-RNA was monitored by analysis on 6% polyacrylamide-8M urea gels, where the amounts of full-length products were generally ≥ 90%.

Radioactive microRNA anti-sense probes were constructed using ‘mirVana™ Probe Construction Kit (Thermo Scientific) according to manufacturer’s instructions.

### Northern Blot Analysis

Total cellular RNA was extracted from DMSO- and WA-treated MCF-7 cells according to the protocol described previously [62].

### RNA affinity column chromatography

*In vitro*-transcribed polyadenylated *bec-A* RNA [using a poly A kit (Ambion) according to the manufacturer’s protocol] was incubated with the oligo (dT)-agarose beads (100 μg in 200 μl) for 2 h at 4°C, then incubated with the samples (pre-cleared cytosolic extract of WA-treated MCF-7 cells) for 2 h at 4°C with gentle shaking followed by centrifugation at 10,000 *r.c.f*. for 1 min. The proteins were eluted from the matrix with a step gradient of NaCl (0.2 to 1.0 M; 0.2 M steps). Eluted fractions were run on 10% SDS-PAGE.

### RNA-protein co-immunoprecipitation assay

Immunoprecipitation of RNA-protein complexes was performed as described in [29]. Briefly, WA-treated MCF-7 cells (5×10^7^) in PBS were cross-linked with formaldehyde [final concentration of 0.1% (v/v)], then quenched with glycine (pH 7.0, 0.25 M). The pre-cleared cell extracts were incubated for 2 h with shaking at 4°C with 20µl protein A/G sepharose beads which were pre-incubated for 1 h with 4μg of monoclonal anti-HuR, anti-GW-182, anti-Ago2 and normal mouse IgG (0.2 μg/µl) antibodies. The A/G sepharose beads were washed, reverse cross-linked at 70°C for 45 mins. Finally, RNA was extracted from the immunoprecipitated samples using TRIZOL, treated with DNase I, reverse transcribed and amplified by semi-quantitative PCR for observing mRNA levels of *becn1* and β*-actin*. Results were also quantified by qRT-PCR.

### Polysome Analysis

MCF-7 cells (WA-treated or transfected) were homogenized in polysome lysis buffer containing cycloheximide (0.1 mg/ml). Preparation of 10-50% (w/v) sucrose gradient and collection of fractions was done as cited in [30]. RNA was isolated from individual fractions by phenol-chloroform extraction followed by ethanol precipitation while proteins from individual fractions were precipitated with 30% tri-chloro acetic acid (TCA) followed by acetone wash and re-suspension in 1X protein loading buffer [30].

### Statistical Analysis

Parametric unpaired *t*-test was used for analysis of statistical significance of graphs. *P*-values <0.05 were considered to be statistically significant while P>0.05 were considered non-significant (NS).

## Supplementary Figure legends

**Figure S1. Translational yield of beclin-1 in WA-treated MCF-7 cells.**

Translational yield (TY) was calculated by measuring the ratio of protein levels (western blot band intensity of protein level of Beclin1 *w.r.t.* β-Actin) with their corresponding mRNA levels (measured by qRT-PCR: mRNA level of *becn1 w.r.t. β-actin*) for control and 24 hrs WA-treated sample.

**Figure S2. Expression of Ago2 protein in WA-treated MCF-7 cells.**

(S1.i) Western blots of Ago2 with MCF-7 cells DMSO- or WA-treated,with whole cell extracts where β-Actin was used as loading control. 80μg of total proteins were resolved in 8% SDS-PAGE. (S1.ii) Immunofluorescence analysis was done by confocal microscopy (magnification 100X) to check cellular localization of Ago2 (using FITC-conjugated antibody; green) where nucleus was stained with DAPI (blue).

## Acknowledgment

The authors wish to thank Dr. ParthoSarothi Ray and Prof. Tapas Sengupta (IISER, Kolkata, India) for using the ultra-centrifuge and polysome fractionators and their valuable thoughts. We also like to thank Ms. Monalisa Mondal for helping us handling of ultra-centrifugation and polysome fractionation. We acknowledge CSIR forproviding the fellowship to S De, S Das andUGC for providing the fellowship to AC, SG. We acknowledge DST-FIST and UGC-CAS program for providing funds and instrumental facilities in our department.

## Data availability

All relevant data supporting the key findings of this study are available within the article and the Supplementary files.

## Abbreviations

becn1: beclin1
mRNA: messenger RNA
UTR: untranslated region
ARE: AU-rich element
RNABP: RNA binding protein
HuR: Human Antigen R; small interfering RNA
SGs: Stress Granules
miRNA: microRNA
RISC: RNA induced silencing complex
GW-bodies: glycine-tryptophan protein of 182 kDa.

