## Supplementary figures and images for "Interplay of microRNA-20a and HuR in the regulation of beclin1 during Withaferin-A mediated impaired autophagy in Breast Cancer cell-line, MCF-7"

### Supplementary figures

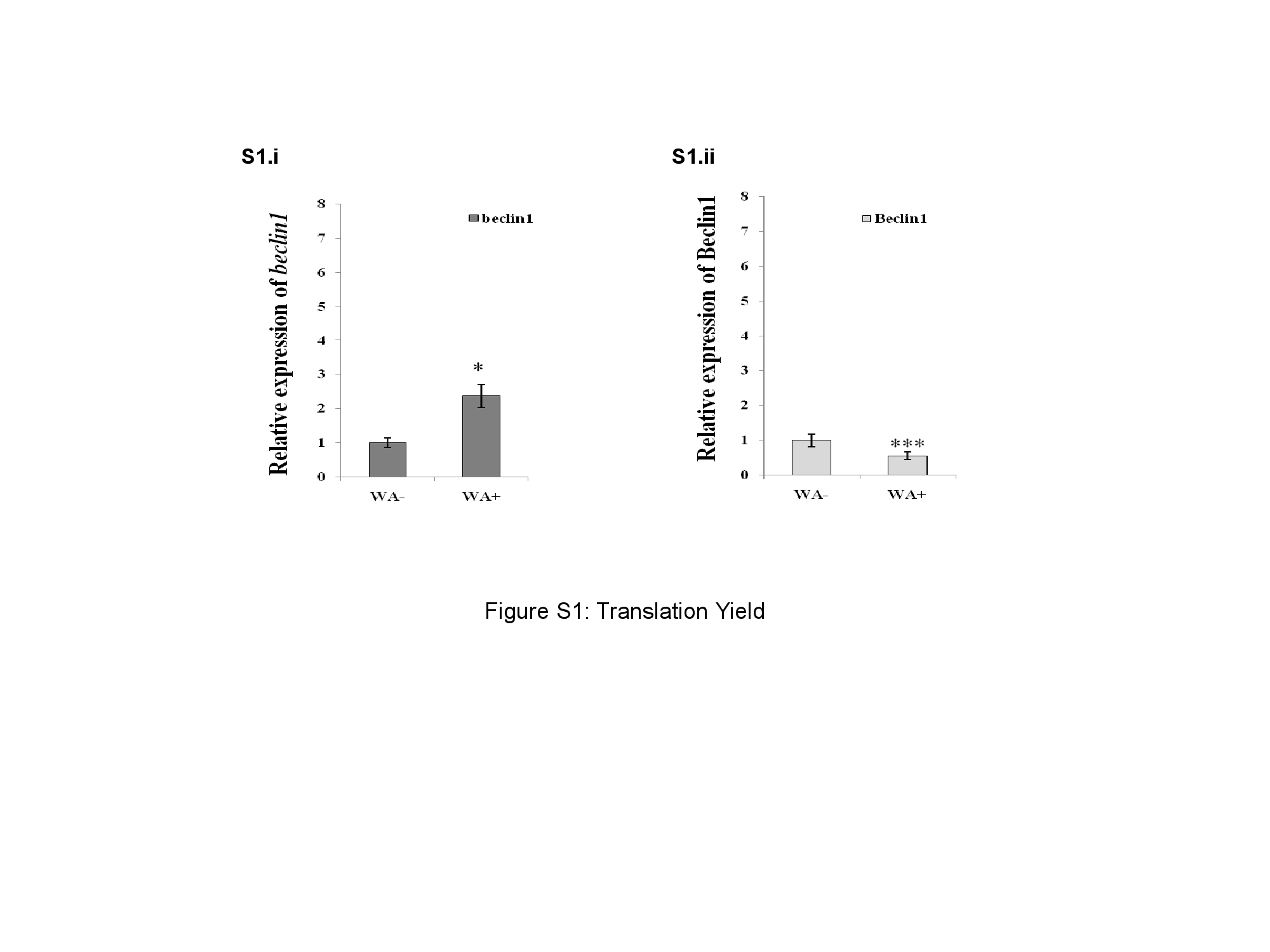

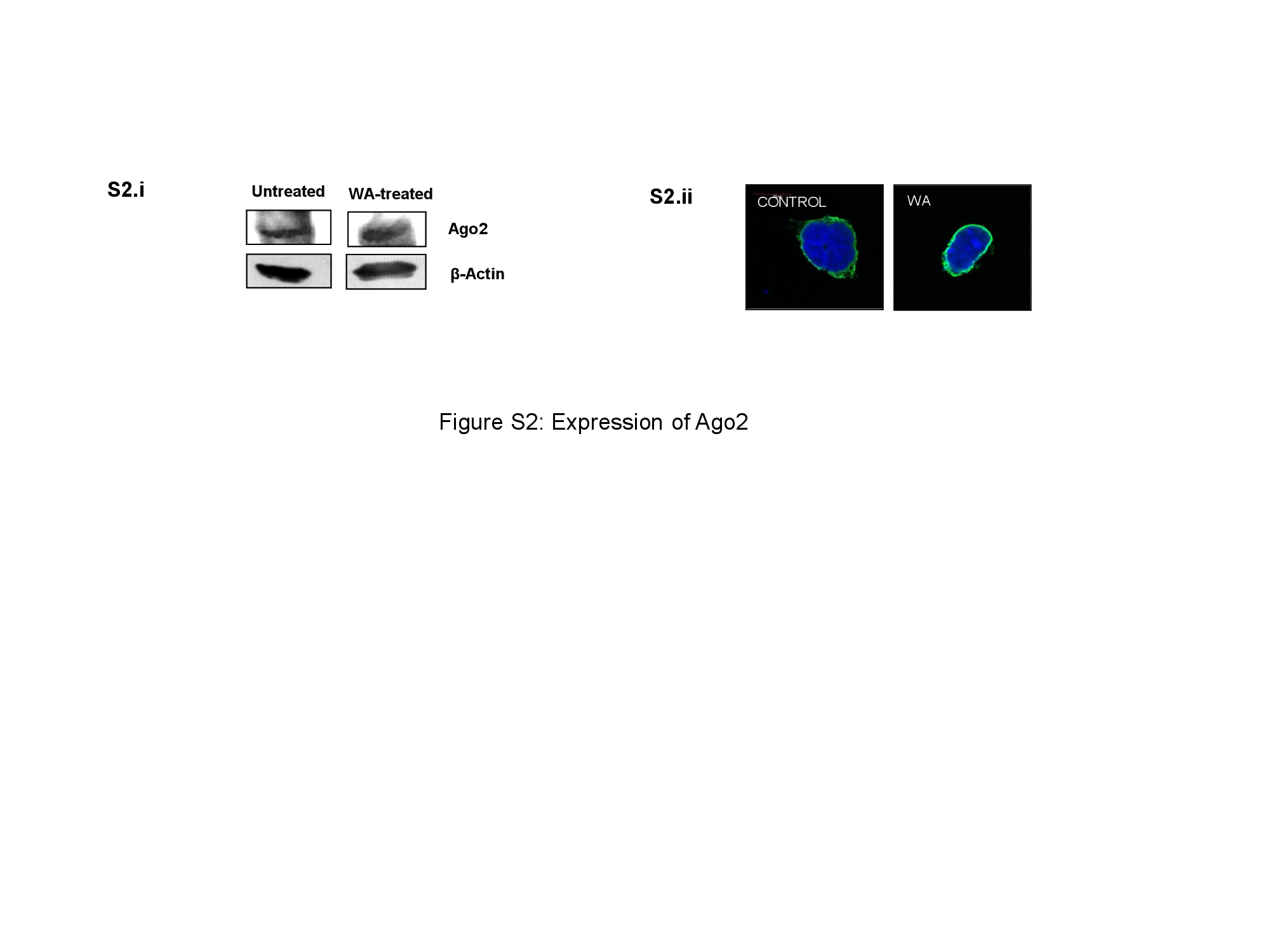
